# Characterizing dynamic neural representations of scene attractiveness

**DOI:** 10.1101/2022.04.27.489648

**Authors:** Daniel Kaiser

**Affiliations:** Mathematical Institute, Department of Mathematics and Computer Science, Physics, Geography, Justus-Liebig-University Gießen, Germany, Center for Mind, Brain and Behavior (CMBB), Philipps-University Marburg and Justus-Liebig-University Gießen, Germany

## Abstract

Aesthetic experiences during natural vision are varied: they can arise from viewing scenic landscapes, interesting architecture, or attractive people. Recent research in the field of neuroaesthetics has taught us a lot about where in the brain such aesthetic experiences are represented. Much less is known about when such experiences arise during the cortical processing cascade. Particularly, the dynamic neural representation of perceived attractiveness for rich natural scenes is not well understood. Here, I present data from an EEG experiment, in which participants provided attractiveness judgments for a set of diverse natural scenes. Using multivariate pattern analysis, I demonstrate that scene attractiveness is mirrored in early brain signals that arise within 200ms of vision, suggesting that the aesthetic appeal of scenes is first resolved during perceptual processing. In more detailed analyses, I show that even such early neural correlates of scene attractiveness are partly related to inter-individual variation in aesthetic preferences and that they generalize across scene contents. Together, these results characterize the time-resolved neural dynamics that give rise to aesthetic experiences in complex natural environments.

## Introduction

In our daily lives, aesthetic experiences can arise from a variety of different visual contents, such as from a scenic sunset, impressive historical architecture, or an attractive face. In the field of neuroaesthetics, one of the key questions is how the aesthetic appeal of such diverse experiences is dynamically extracted by the brain (Pearce et al., 2016; Skov & Nadal, 2020).

Over the recent decades, we have learned a lot about *where* in the brain aesthetic experiences are processes. Researchers have used fMRI to pinpoint the neural correlates of perceived aesthetic appeal for a range of stimuli, such as faces, landscapes, abstract patterns, and artworks (Isik & Vessel 2021; Jacobsen et al., 2006; Kawabata & Zeki, 2004; Pegors et al., 2015; Vessel et al., 2019; Winston et al., 2007; Yue et al., 2006; Zhao et al., 2020). Some of these studies also demonstrate that the aesthetic appeal of vastly different visual contents, such as faces and scenes (Pegors et al., 2015) or artworks and photographs (Vessel et al., 2019) is represented in similar regions across cortex, including parts of the visual cortex and the frontal cortex, as well as the default mode network. These studies suggest that aesthetic experiences can not only arise from seemingly dissimilar visual stimuli, but that these experiences yield similar cortical correlates.

Much less is known about *when* aesthetic experiences arise dynamically across the cortical processing cascade. Much of the M/EEG literature addressing this question has focused on the perception of face attractiveness (Carbon et al., 2018; Kaiser & Nyga, 2020; Schacht et al., 2008; Werheid et al., 2007; Zhang & Deng, 2012), with many studies highlighting that a face’s attractiveness can impact early and fundamental stages of the face processing hierarchy. In our own work, we have further highlighted that such early representations of face attractiveness are partly explained by personal preferences, rather than only by attractiveness judgments that are shared among a large group of observers (Kaiser & Nyga, 2020), suggesting that even the perceptual correlates of attractiveness are shaped in personally idiosyncratic ways. Unlike the rich literature on face attractiveness, only few studies have looked at the time-resolved neural correlates of aesthetic judgments for other stimuli, such as abstract patterns (Höfel & Jacobsen, 2007; Jacobsen & Höfel, 2003) and various types of artworks (Cela-Conde et al., 2004; de Tommaso et al., 2007; Strijbosch et al., 2021). However, we currently do now know how the brain dynamically represents attractiveness for the natural scenes we typically experience during our everyday lives.

There are three open questions about the neural dynamics underlying perceived scene attractiveness: (1) Is the aesthetic appeal of natural scenes already represented during early stages of cortical processing, indicating that differences in fundamental perceptual processes are involved in the extraction of scene attractiveness? (2) Are judgments of scene attractiveness based on a general consensus of what is aesthetically pleasing or are they partly shaped by personal aesthetic preferences? (3) Is the aesthetic appeal of natural scenes extracted similarly across different scene contents?

To answer these questions, I investigated how the perceived aesthetic appeal of a set of 100 diverse natural scene images is mirrored in temporally resolved EEG signals. For this, I use a multivariate pattern analysis framework that we recently developed in an EEG study of face attractiveness (Kaiser & Nyga, 2020). Specifically, I modelled how the pairwise similarity between scenes in the EEG signals over time was related to the similarity in aesthetic appeal ratings. The resulting data demonstrate that (1) the aesthetic appeal of scenes is encoded in early brain signals emerging within 200ms after scene presentation, (2) part of this early neural correlate of scene attractiveness is explained by inter-individual variation in aesthetic perception, and (3) even early neural correlates of perceived aesthetic appeal generalize across a variety of scene contents.

## Materials and Methods

### Participants

24 healthy adult participants took part (mean age 19.6 years, SD=1.7; 21 female). This sample size was identical to our previous EEG study investigating face attractiveness (Kaiser & Nyga, 2020). One participant was excluded due to a technical error in the recordings, leaving a final sample of 23 participants. Participants received course credits. All participants provided written informed consent. Procedures were approved by the ethical committee of the Department of Psychology, University of York, and were in accordance with the Declaration of Helsinki.

### Stimuli

The stimulus set consisted of 100 natural scene photographs. All stimuli were resized to 600-by-400 pixels. The stimuli were chosen from the top- and bottom-rated images contained in the AVA (Murray et al., 2012) and photo.net (Datta et al., 2008) aesthetic rating databases. To match the overall visual content across images that were expected to yield high or low aesthetic appeal ratings, stimuli were arranged into pairs (Fig. 1a): in each pair, the two images depicted visually and/or conceptually similar contents (e.g., a group of people or an indoor room). To check whether there were any consistent visual differences across pairs, I computed GIST descriptors (Oliva & Torralba, 2001) for all images. I then checked whether the high attractiveness and low attractiveness images within each pair were reliably different. To this end, I trained a nearest-neighbor classifier (as implemented in CoSMoMVPA; Oosterhof et al., 2016) to classify the images’ attractiveness into high and low. This classifier was trained on the GIST descriptors for all but one pairs and then tested on the remaining pair of images (this procedure was repeated for each pair being left out). The GIST-based classifier could not successfully discriminate between the images that were taken from the top- and bottom-rated images in the database (49% correct). Similar results were obtained using the “pooling” layers of an HMAX (Riesenhuber & Poggio, 1999; Serre et al., 2007) filter-bank model (layer C1: 45% correct; layer C2: 49% correct). The pairing of stimuli thus ensured that there were no pronounced visual differences in basic visual characteristics between images of different aesthetic appeal (as indicated by their database rankings). In all subsequent analyses, I focused on the ratings provided by participants in the current study.

**Figure 1.**
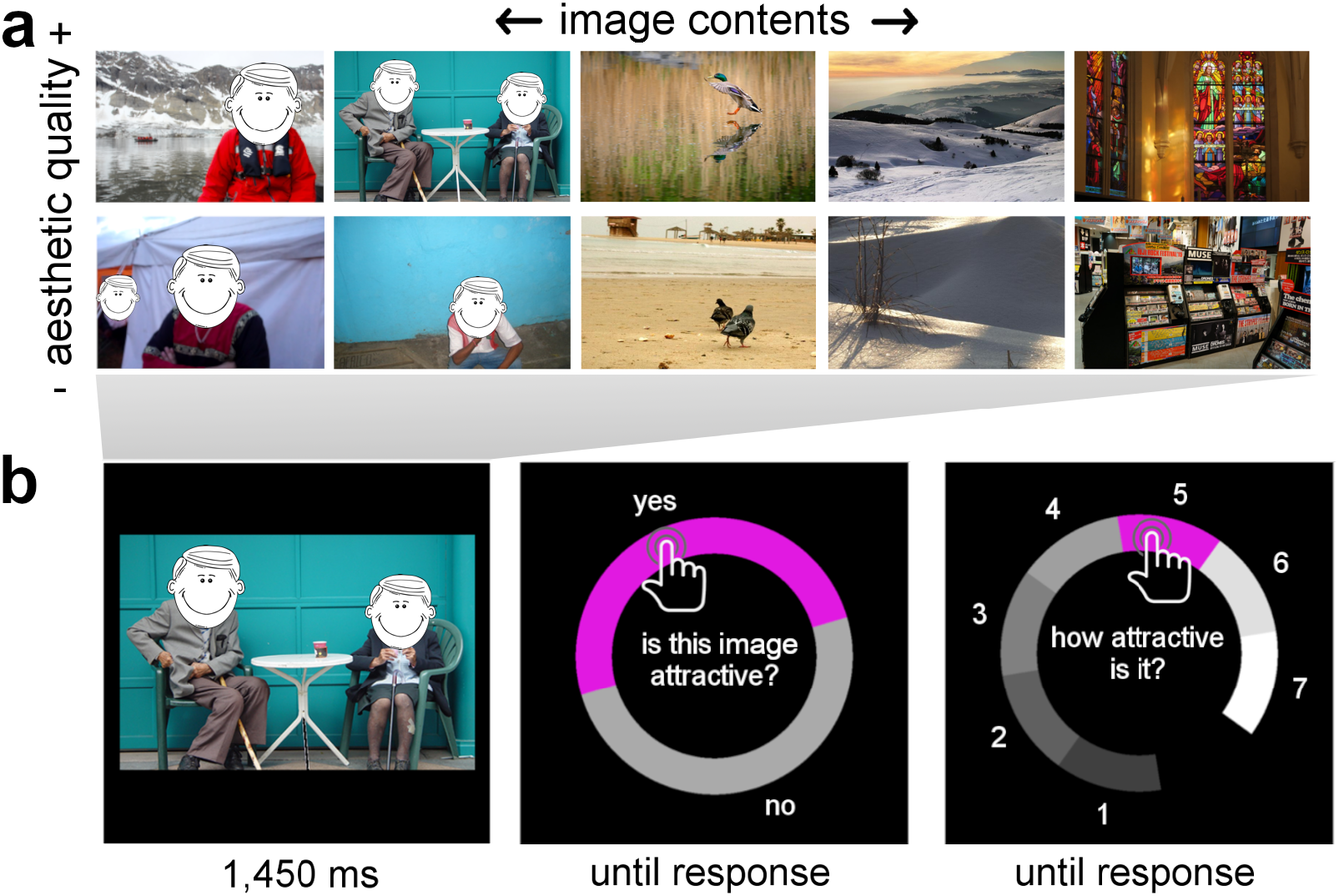
Stimuli and Paradigm. **a)** The stimulus set consisted of 100 natural scene photographs depicting a variety of diverse contents. Stimuli were arranged into visually similar pairs of high and low aesthetic appeal (based on visual aesthetics databases) to reduce the visual variability between scenes that were likely judged as relative attractive or unattractive. **b)** During each trial of the EEG experiment, participants viewed a single scene and subsequently rated its attractiveness on two complementary response screens asking whether the image was attractive (“yes/no response”) and how attractive it was on a 1-7 scale (“rating response”). Response options were arranged randomly around a circular response screen to prevent motor preparation during the image presentation. Trials were separated by an 800-1200ms inter-trial interval.

### Paradigm

The experimental paradigm and analysis approach was largely identical to our recent EEG study on face attractiveness perception (Kaiser & Nyga, 2020). Participants viewed a single scene on every trial (8 by 5.3 degrees visual angle), which was presented for 1,450ms on a uniform black background (Fig. 1b). After a 100ms blank screen, participants were asked to give two complementary responses: First, they were asked to provide a binary attractiveness response, indicating whether they found the scene attractive or not (hereinafter referred to as “yes/no response”). Second, they were asked to provide a more fine-grained rating, indicating how attractive they found the scene on a 1 to 7 rating scale (hereinafter referred to as “rating response”). Both ratings were given with the mouse and were non-speeded. Participants were instructed to indicate, how aesthetically pleasing, or attractive, or beautiful they rated each image (without any distinction between these concepts). To prevent participants from preparing a motor response, the response options for both responses were presented at random angular positions across a circular response screen (Fig. 1b). Participants were further instructed to keep central fixation on a pink fixation dot during the scene presentation and to restrict eye blinks to the period when they selected their responses. Trials were separated by an inter-trial interval randomly varying between 800ms and 1,200ms. The experiment consisted of 7 blocks, in each of which each image was shown once, in random order. Each image was therefore repeated 7 times, yielding 700 trials in total. The experiment was carried out in a dimly lit and quiet room. Stimuli were presented on a VIEWPixx display with a 1920-by-1020 resolution and stimulus presentation was controlled using the Psychtoolbox (Brainard, 1997).

### EEG acquisition and preprocessing

EEG signals were recorded using an ANT Waveguard 64-electrode system and a TMSi REFA amplifier. Electrodes were arranged in accordance with the standard 10–10 system. EEG data were recorded at 250Hz sampling rate using the ANT Neuroscan Sofware. Offline preprocessing was performed using FieldTrip (Oostenveld et al., 2011). EEG data were referenced to the Fz electrode (which was discarded after preprocessing), epoched from – 500ms to 1900ms relative to stimulus onset, and baseline-corrected by subtracting the mean pre-stimulus signal for each electrode. A band-pass filter was applied to remove 50Hz line noise. Channels and trials containing excessive noise were removed based on visual inspection. On average, 9 channels (SE=0.6) and 69 trials (SE=11) were removed. Blinks and eye movement artifacts were removed using independent component analysis and visual inspection of the resulting components. After preprocessing, EEG epochs were cropped from −250ms pre-stimulus to 1450ms poststimulus.

### Extracting neural representational similarity

To track representations across time, I used representational similarity analysis (RSA; Kriegeskorte et al., 2008). First, neural RDMs were constructed separately for each participant, using the CoSMoMVPA toolbox (Oosterhof et al., 2016). RDMs were created for 34 consecutive time bins of 50ms width, from −250ms to 1450ms relative to scene onset. The following analyses were done separately for each time bin. At each bin, response patterns were extracted across 12 time points (covering 50ms at 250Hz) and 63 electrodes (after preprocessing, electrode counts could be lower for individual participants). These data were then unfolded into a 756-element vector. Before RDM construction, I performed principal-component analyses (PCAs) to reduce the dimensionality of the response vectors (Grootswagers et al., 2017; Kaiser et al., 2020a). I split the available data into two independent subsets, with an equal number of trials per condition randomly assigned to each subset. The first subset of the data was used to perform the PCA decomposition. The PCA decomposition was then projected onto the second subset, retaining only the components needed to explain 99% of the variance in the first subset (97 components on average, SD across time: 19, SD across participants: 16). RDMs were constructed from the second subset. I first averaged across all available trials for each condition, and then correlated (Spearman-correlations) the response vectors for each pairwise combination of scenes. These correlations were subtracted from 1 and arranged into a 100-by-100 RDM. Each entry in this RDM reflected a measure of neural dissimilarity for a specific pair of scenes. RDM diagonals were always empty. This procedure was then repeated with the two subsets swapped. Finally, the whole analysis was repeated 50 times, with trials assigned randomly to the two subsets each time. RDMs were averaged across all repetitions, yielding a single RDM for each time bin.

### Modelling representational similarity

Neural RDMs were modelled by a set of predictor RDMs that also spanned 100 by 100 entries and captured the scenes’ pairwise similarities on a set of candidate properties. Correspondence between the predictor RDMs and the neural RDMs was assessed by correlating (Spearman-correlations) all off-diagonal entries between the two types of RDMs, separately for each time point at which a neural RDM was available, yielding a time course of correspondence. I performed three types of RSA, using three types of predictor RDMs: First, to model how well neural representations were predicted by attractiveness judgments, I created two predictor RDMs that captured the scenes’ similarities in the yes/no responses and the rating responses. In these RDMs, each entry was computed by taking the absolute difference between the average yes/no responses (coded as 2:yes and 1:no) or attractiveness ratings between two scene images. These RDMs were constructed separately for each participant. Second, I tested whether the complex feature organization emerging in a deep neural network (DNN) model of categorization explains the correspondence between attractiveness judgments and neural representation. Although the stimulus set was matched in approximate content and low-level features across the more or less attractive images, the images may still differ in a set of complex visual features that is routinely read out during visual categorization: to quantify the extraction of such features, I employed a DNN as a model of the cortical processing hierarchy (Cichy & Kaiser, 2019; Kriegeskorte, 2015). I extracted features along the 16 layers of a VGG16 DNN (Simonyan & Zisserman, 2014) trained on ImageNet (Deng et al., 2009). For each layer, I then constructed an RDM by computing 1 minus the correlation of activation vectors for each pair of scenes. RDMs were then averaged within 6 layers blocks along the network (5 blocks of convolutional layers and 1 block of fully connected layers; see Fig. 4a). These RDMs were then correlated with the neural RDMs to determine how well the DNN predicted the neural data in the first place. After that, I conducted partial correlation analyses in which I re-performed the correlation between the aesthetic judgment RDMs and the neural RDMs while partialing out the RDMs for each DNN layer block. Third, to model how well average and individual attractiveness judgments predict the neural data, I created RDMs that captured the scenes’ similarities in the average yes/no responses and the rating responses across a set of participants. These were created in the same way as the previous RDMs, but for each participant, I created an average RDM that was based on the average yes/no responses or rating responses from all other participants. In a partial correlation analysis, I then again correlated the predictor RDMs for the yes/no responses and ratings responses with the neural RDMs, but now partialing out the average response across all other participants. This yielded an estimate of how well individual responses account for cortical representations when average responses are controlled for (Kaiser & Nyga, 2020). Fourth, to test whether the neural correlates of scene attractiveness generalize across scene categories, I performed analyses in which the correspondence between the predictor RDMs for the yes/no responses or rating responses and the neural RDMs was only assessed for those pairwise comparisons in which the two scenes came from opposite categories. This analyses thus reveals whether attractive scenes from one category are represented similarly as attractive scenes from another category (and dissimilarly from less attractive scenes from another category), thus revealing a relatively content-independent representation of aesthetic appeal. I tested such generalization for three categorical distinctions: scenes containing (n=42) versus not containing (n=58) people, natural (n=58) versus man-made (n=42) scenes, and scenes containing (n=38) versus not containing (n=62) prominent foreground objects. Scenes were assigned to these labels by the author.

**Figure 2.**
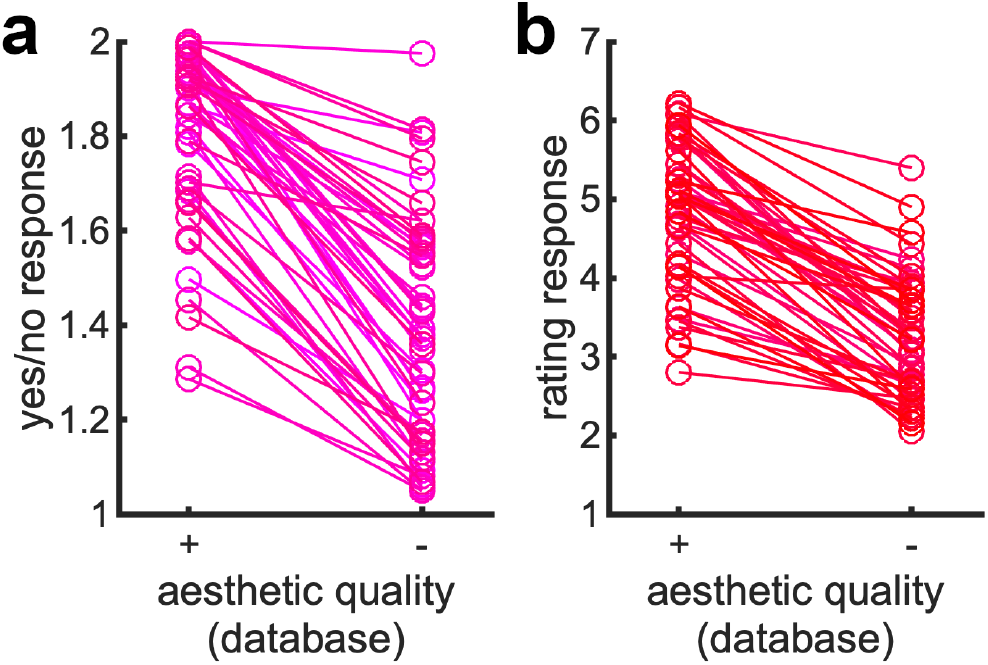
Analysis of attractiveness judgments. **a)** Average yes/no responses across stimulus pairs (2:yes, 1:no; each line represents a pair). **b)** Average rating responses across stimulus pairs (7:best, 1:worst; each line represents a pair). For both judgments and within each pair, the scene that had the higher attractiveness rating in the aesthetics databases also received the higher yes/no response and rating response in the experiment.

**Figure 3.**
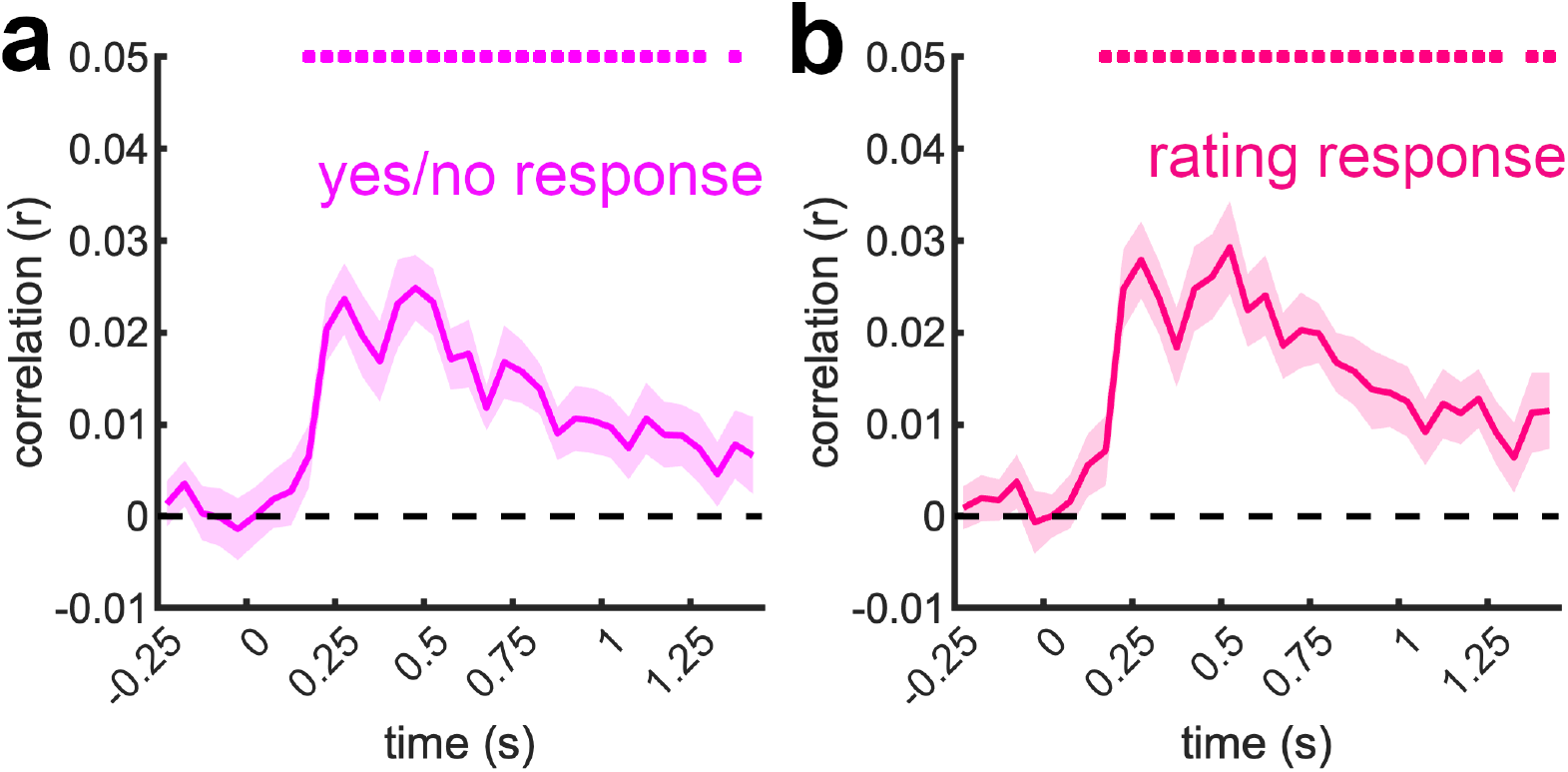
Neural dynamics of perceived scene attractiveness. **a)** Average correlations between the neural RDMs (reflecting pairwise similarities between scenes at each time point) and a predictor RDM based on participants’ yes/no responses (reflecting pairwise similarities in how similarly scenes were judged as attractive or unattractive). **b)** Average correlations between the neural RDMs and a predictor RDM based on rating responses (reflecting pairwise similarities in how similarly scenes were rated). Both types of attractiveness judgments predicted neural responses from the 150-200ms time bin and in a temporally sustained fashion. Error margins represent standard errors of the mean and significance markers denote p_corr_<0.05.

**Figure 4.**
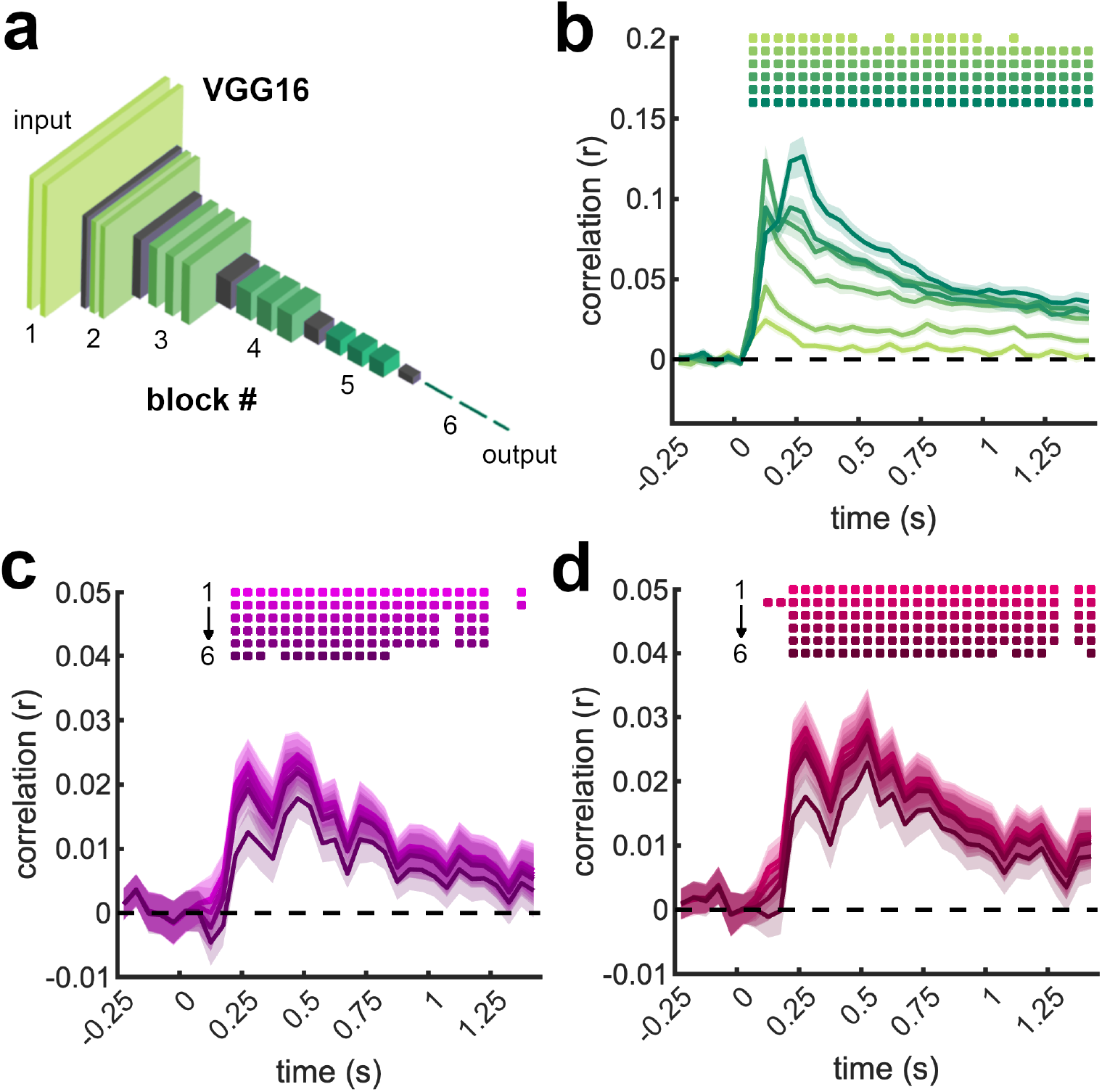
Controlling for DNN features. **a)** To approximate the extraction of categorization-related features in the visual system, I used a VGG16 DNN trained on ImageNet. From this network, I extracted RDMs for 6 layer blocks along the network, as illustrated; the last three layers are the network’s fully connected layers. **b)** The RDMs extracted from the DNN predicted neural activations well, with early layers predominantly predicting relatively early activations and late layers offering more accurate prediction of later activations. **c)** Re-performing the correlations between the RDMs based on yes/no attractiveness responses and the neural RDMs (as in Fig. 3a) while partialing out the DNN RDMs obtained from different layer blocks showed that DNN features at different depths cannot account for the correspondence between attractiveness judgments and brain representations. **d)** Similar results were obtained for the rating responses. Error margins represent standard errors of the mean and significance markers denote p_corr_<0.05.

### Statistical testing

Correlations between neural and predictor RDMs were compared to zero using one-sample t-tests (one-sided against zero) for each time bin. The resulting p-values were corrected for multiple comparisons using FDR-corrections. Test statistics (t-values) and effect sizes (Cohen’s d) are provided for all peak effects.

### Data Availability

All data are available on OSF (doi.org/10.17605/OSF.IO/SKHF7). Other materials will be made available upon request.

## Results

### Ratings of scene attractiveness

The behavioral responses given during the experiment confirmed the pairing of images in our stimulus set, with stimuli obtained from the top-rated database images yielding consistently higher perceived aesthetic appeal than those obtained from the bottom-rated database images. Indeed, the expected difference within each pair was consistent across all pairs and both for the yes/no responses (Fig. 2a) and the rating responses (Fig. 2b). Across participants, the top-rated images yielded significantly higher yes/no responses (t[22]=12.6, p<0.001) and rating responses (t[22]=11.7, p<0.001) then the bottom-rated images.

### Neural dynamics of perceived scene attractiveness

To track the emergence of neural responses related to scene attractiveness, I correlated neural RDMs with predictor RDMs that captured the images’ pairwise similarities in participants’ yes/no and rating responses. Yes/no responses (Fig. 3a) significantly predicted cortical responses from the 150-200ms time bin (peaking at 450-500ms, peak t[22]=6.94, p_corr_<0.001, d=1.45). Rating responses (Fig. 3b) also predicted cortical responses from the 150-200ms time bin (peaking at 500-550ms, peak t[22]=5.80, p_corr_<0.001, d=1.21). This shows that the aesthetic appeal of natural scenes is rapidly and sustainedly contained in EEG signals, suggesting that aesthetic experiences manifest during early stages of visual information processing and are encoded in brain responses for extended periods of time.

### Controlling for DNN features

Although the more and less attractive images in the stimulus set were matched for their approximate visual content and for low-level features, the correspondence between aesthetic judgments and neural representations may still be accounted for when more complex visual features are considered. To test this possibility, I compared neural representations to the representations emerging in a DNN model of visual categorization (VGG16; Simonyan & Zisserman, 2014; see Figure 4a). When correlating RDMs extracted from 6 layer blocks along this DNN with the neural RDMs, I found strong signatures of hierarchical correspondence (Cichy et al., 2016), with early layers predominantly predicting early brain activations, and late layers more strongly predicting later activations (Fig. 4b). This result shows that the DNN was able to approximate visual feature processing in cortex. To test whether such categorization-related feature processing could account for the correspondence between attractiveness judgments and neural representations, I correlated the RDMs constructed from the yes/no responses and rating responses with the neural RDMs while partialing out the RDMs obtained from the DNN layer blocks. Across the board and for both types of responses, correlations remained significant (Fig. 4c/d), indicating that the features extracted during categorization (as approximated by the DNN) are not sufficient for explaining the cortical representation of scene attractiveness. Partialing out the RDMs from the last layer block (i.e., the fully connected layers of VGG16) produced the greatest drop in correlations, with a significant reduction compared to the original results, both for the yes/no responses and the rating responses (from 50-100ms and across the whole epoch). However, correlations remained well above chance, both for the yes/no responses (from the 200-250ms time bin, peaking at 450-500ms, peak t[22]=5.69, p_corr_<0.001, d=1.19) and the rating responses (from the 200-250ms time bin, peaking at 500-550ms, peak t[22]=4.90, p_corr_<0.001, d=1.02). This indicates that aesthetic appeal is not solely a reflection of features that are read out during categorization, but that qualitatively different features determine the attractiveness of natural visual inputs.

### Personal versus shared aesthetic judgments

Next, I investigated whether personal ratings of aesthetic appeal predict cortical responses beyond the average ratings provided by a group of observers (Kaiser & Nyga, 2020). I thus correlated the neural RDMs with additional predictor RDMs that for each participant were constructed from the average responses of all other participants in the experiment. For the yes/no responses (Fig. 5a), averaged responses from all other participants predicted neural responses as well as participants’ own responses, starting from the 200-250ms time bin (peaking at 250-300ms, peak t[22]=6.47, p_corr_<0.001, d=1.35). A similar pattern emerged for the rating responses (Fig. 5b), starting from the 200-250ms time bin (peaking at 250-300ms, peak t[22]=6.76, p_corr_<0.001, d=1.41). For both measures, there were no significant differences between participants’ own ratings and the average of others’ ratings (all p_corr_>0.93). Given this similarity, I next tested whether other people’s ratings explain the correspondence between participants’ own ratings and the neural responses. I thus performed a partial correlation analysis, correlating the neural RDMs with the RDMs constructed from participants’ own responses, partialing out the average responses from all other participants. This analysis still revealed a significant correlation, both for the yes/no responses (from the 200-250ms time bin, peaking at 400-450ms, peak t[22]=3.97, p_corr_=0.004, d=0.83) and the rating responses (from the 150-200ms time bin, peaking at 500-550ms, peak t[22]=5.16, p_corr_<0.001, d=1.08). This finding suggests that although both personal and shared ratings can predict (early) neural representations, these ratings explain complimentary shares of the neural dynamics: If predictions based on average judgments captured essentially identical variance in the neural data as predictions based on individual judgments, partialing out the RDM based on average judgments should have eliminated the correlation between the RDM based on individual judgments and the neural data. That individual judgments continued to predict neural representations in this analysis thus suggests that brain representations of aesthetic appeal cannot be fully accounted for by a form of “average taste”, but that individual ratings of aesthetic appeal are needed for more accurate prediction.

**Figure 5.**
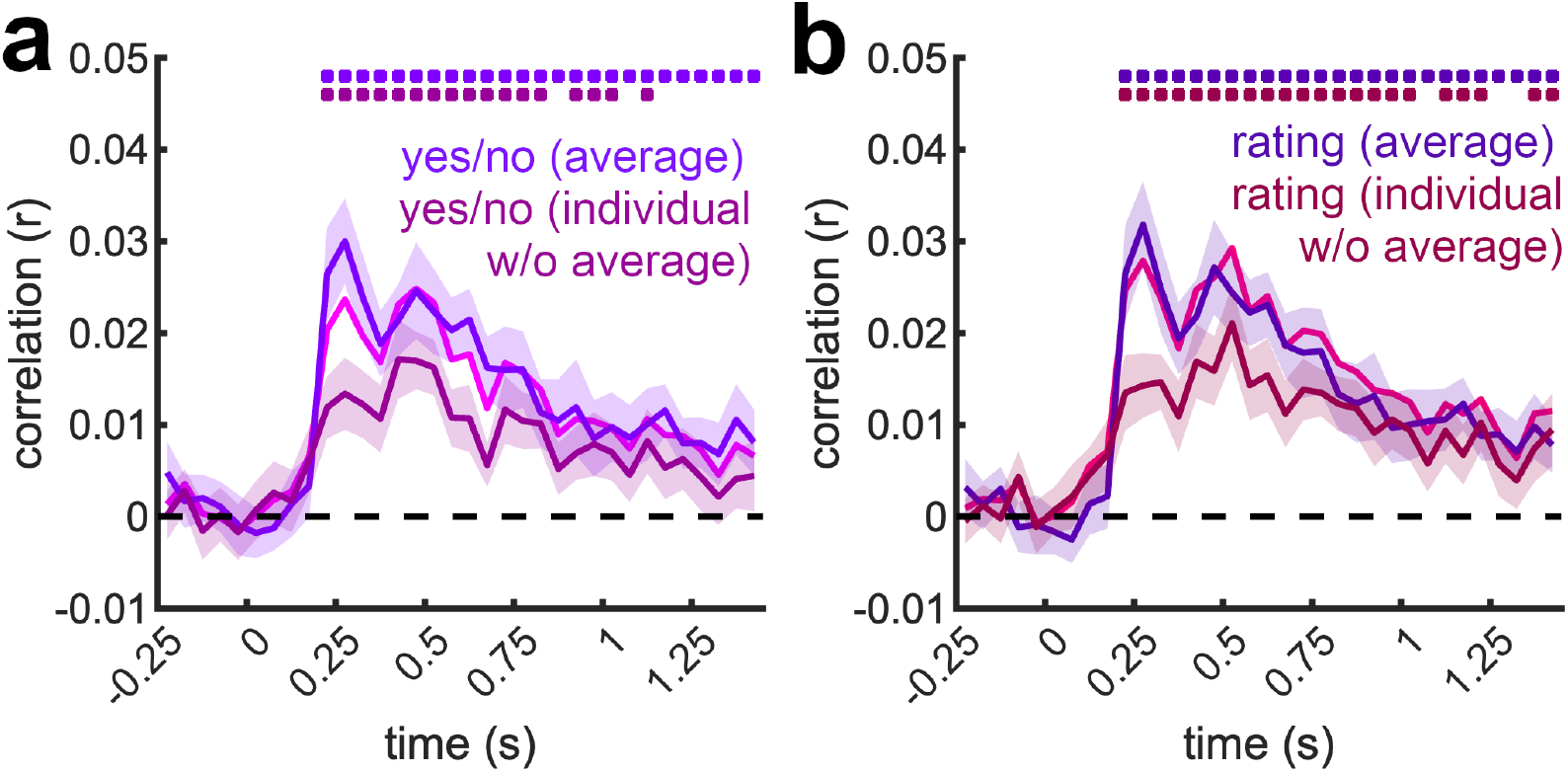
Neural representation of personal aesthetic judgments. **a)** Correlations between the neural RDMs and a predictor RDM based on the average yes/no responses of all other participants modelled the data equally well as the predictor RDM based on each individual participant’s yes/no responses. Results from the previous analysis (pink line) are shown for reference (same as in Fig. 3a). The individual yes/no responses still predicted neural responses well when the average yes/no response across all people was controlled for, suggesting that inter-individual variability in aesthetic judgments is reflected in (early) cortical responses. **b)** A highly similar pattern of results emerged when the rating responses were analyzed. Results from the previous analysis (red line) are again shown for reference (same as in Fig. 3b). Error margins represent standard errors of the mean and significance markers denote p_corr_<0.05.

### Extracting aesthetic appeal across scene contents

A key question in the literature is whether the aesthetic appeal of different content types is extracted through similar or different cortical mechanisms. However, a particular focus in the literature has thus far been placed on the attractiveness of humans. In a first analysis, I therefore asked whether aesthetic appeal is extracted differently for scenes that depict humans and scenes that do not. For this purpose, I split the stimulus set into scenes that did or did not depict humans. I then correlated the neural RDMs and predictor RDMs, only using the pairwise comparisons across scenes with and without humans (Fig. 6a). This analysis revealed a significant correspondence between perceived aesthetic appeal and brain responses, both for the yes/no response (from the 200-250ms time bin, peaking at 450-500ms, peak t[22]=6.22, p_corr_<0.001, d=1.30) and the rating response (from the 200-250ms time bin, peaking at 500-550ms, peak t[22]=5.57, p_corr_<0.001, d=1.16). Second, I asked whether aesthetic appeal is extracted similarly for natural and man-made scenes, as suggested by previous fMRI findings (Vessel et al., 2019). I then again performed the RSA only using pairwise comparisons across the natural and man-made scenes (Fig. 6b). This analysis also revealed a significant correspondence between perceived attractiveness and brain responses, both for the yes/no response (from the 200-250ms time bin, peaking at 200-250ms, peak t[22]=4.30, p_corr_=0.005, d=0.90) and the rating response (from the 200-250ms time bin, peaking at 500-550ms, peak t[22]=4.28, p_corr_=0.002, d=0.89). Third, I asked whether scene attractiveness is represented differently when it is potentially derived primarily from prominent foreground objects (e.g., a person or an animal occupying the scene foreground), rather than more global scene information. I thus tested whether aesthetic appeal is extracted differently for scenes that had a dominant foreground object, which could allow for a rapid aesthetic appraisal solely based on a single object, and scenes that did not. I then performed the RSA only using pairwise comparisons across the foreground-object and no-foreground-object scenes (Fig. 6c). Yet again, this analysis revealed a significant correspondence between perceived aesthetic appeal and brain responses, both for the yes/no response (from the 200-250ms time bin, peaking at 450-500ms, peak t[22]=5.92, p_corr_<0.001, d=1.24) and the rating response (from the 150-200ms time bin, peaking at 250-300ms, peak t[22]=5.96, p_corr_<0.001, d=1.24). Together, these analyses show that the early extraction of aesthetic appeal for natural scenes is similar across broad natural contents, suggesting a common neural correlate for aesthetic perception that is already evident at early stages of visual processing.

**Figure 6.**
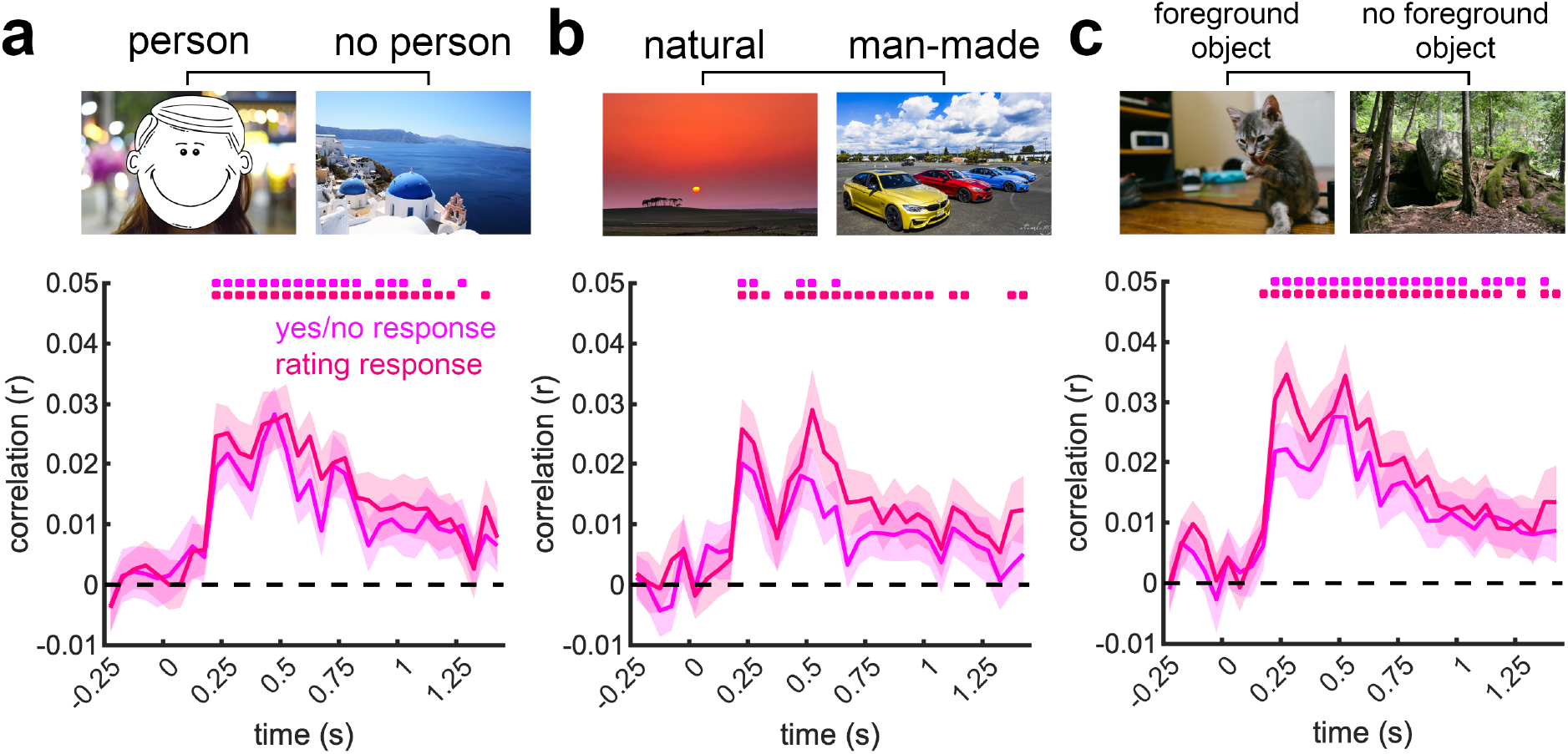
Neural representation of attractiveness across scene contents. **a)** Correlations between the neural RDMs and the predictor RDMs based on yes/no responses or ratings responses when only pairwise comparisons between scenes that did or did not contain a person were considered. **b)** Correlations between the neural RDMs and the predictor RDMs when only pairwise comparisons between natural and man-made scenes were considered. **c)** Correlations between the neural RDMs and the predictor RDMs when only pairwise comparisons between scenes that did or did not contain prominent foreground objects were considered. Together, these three analyses suggest that dynamic neural representations of scene attractiveness are relatively content-independent. Error margins represent standard errors of the mean and significance markers denote p_corr_<0.05.

## Discussion

In this study, I used multivariate pattern analysis on time-resolved EEG data to unveil the neural dynamics of perceived scene attractiveness. I report three key results: (1) Neural representations of scene attractiveness emerge rapidly in brain responses, starting within 200ms of stimulus onset. (2) These early representations are partly explained by inter-individual variation in attractiveness judgments. (3) Neural responses similarly track aesthetic appeal across different scene contents. I will in turn discuss the implications of these three key results.

My findings show that the aesthetic appeal of natural scenes is extracted early on during the visual processing cascade, suggesting that scene attractiveness is partly resolved during perceptual analysis. This is in line with multi-stage models of aesthetic perception that posit an initial sensory processing stage at which aesthetically pleasing features are extracted (Leder et al., 2004; Redies, 2015), as well as with models that explain aesthetic experiences via fluency in sensory processing (Reber et al., 2004). My results further complement EEG work using other visual inputs such as faces (Kaiser & Nyga, 2020; Schacht et al., 2008; Werheid et al., 2007; Zhang & Deng, 2012) or abstract patterns (Jacobsen & Höfel, 2003), which show that aesthetic appeal impacts early visual processing stages. So far, only one other MEG study also included scene photographs as stimuli (Cela-Conde et al., 2004), but primarily found late modulations of cortical responses starting after 400ms. However, most stimuli in this study were artworks, and no separate analyses for artworks and scene photographs were conducted. Together, our study suggests that scene attractiveness is first resolved during fundamental stages of scene analysis, around the time when other scene attributes such as scene category or geometry are analyzed (Cichy et al., 2017; Harel et al., 2016; Kaiser et al., 2020b). Which sensory properties allow scene attractiveness to be resolved during early cortical processing? The matching across relatively attractive and unattractive scenes in the current study suggests that scene attractiveness is not resolved based on a confined set of simple visual features. Further, the failure of categorization-related DNN features in explaining representations of scene attractiveness suggests that the features routinely extracted for the purpose of image categorization are not the same as the features used for determining an images aesthetic appeal. Which features are evaluated for this purpose is a key question for future research. Beyond these early responses, aesthetic appeal was represented in a sustained way. This sustained neural reflection of aesthetic perception may reflect a transitioning from earlier, sensory-driven responses to later, cognitive appraisal processes (Leder et al., 2004; Redies, 2015). The precise nature of this transition, however, needs to be mapped out in future studies that combine temporally resolved EEG recordings with spatially resolved neural recordings.

Interestingly, the current data suggest that the early neural correlates of perceived scene attractiveness are not fully explained by average ratings across participants. When controlling for average attractiveness ratings across participants, brain responses were still predicted by individual participants’ personal ratings. This finding is in line with our previous study on face attractiveness perception (Kaiser & Nyga, 2020), where we could show that early cortical responses are partly predicted by personal attractiveness ratings. Together, these findings suggest that, for multiple visual stimulus categories, even early cortical correlates of aesthetic perception are inherently personal. This notion is not only in line with behavioral studies highlighting substantial inter-individual variation in aesthetic judgment (Hönekopp, 2006; Leder et al., 2016), but also with recent neuroimaging results that show that even basic perceptual representations vary across observers in idiosyncratic ways (Charest et al., 2014). An intriguing possibility that warrants further investigation is that individual differences in the functional architecture of the visual system give rise to different levels of processing fluency (Oppenheimer, 2008; Reber et al., 2004) across stimuli, which ultimately cause idiosyncrasies in aesthetic perception.

My findings further highlight that aesthetic appeal is extracted similarly across different scene contents, such as across (1) scenes that do or do not contain people, (2) natural or man-made scenes, and (3) scene that do or do not contain prominent foreground objects. These results are an important demonstration that even early cortical correlates of perceived aesthetic appeal do not need to be content-specific. They thereby support recent fMRI work that shows a generalization across aesthetic perception for faces, scenes, and artworks (Pegors et al., 2015; Vessel et al., 2019; but see Hu et al., 2020). However, such generalization across contents may ultimately depend on the intrinsic variability in stimulus sets (our stimuli were all rich natural scenes) and the sensitivity of the employed analyses (multivariate pattern analyses may yield higher sensitivity than univariate analyses). Further studies can use the multivariate EEG analysis framework developed here to test generalization across more vastly different visual contents or across modalities.

Together, the current study provides a demonstration for an early cortical correlate of perceived scene attractiveness, which may form the basis of rapid aesthetic evaluation in the wild.

## Acknowledgements

I would like to thank Judit Fiedler and Alex Carter for assisting with EEG data collection. No competing interests are declared.

## Notes

### Competing Interest Statement

The authors have declared no competing interest.

